# Exploring rhythmic and melodic preferences in budgerigars: Individual and possible sex-related variation

**DOI:** 10.64898/2026.08.30.748169

**Authors:** Masumi Wakita

**Affiliations:** Department of Psychology, Teikyo University, 359 Otsuka, Hachioji, Tokyo 192-0395, Japan

**Keywords:** Budgerigar, Preference, Rhythm, Melody, Auditory perception

## Abstract

Budgerigars (Melopsittacus undulatus) are vocal-learning birds with well-developed auditory abilities, but how they behaviorally evaluate melodic and rhythmic structure in sound sequences remains unclear. We examined whether budgerigars show preferences for these acoustic features and whether such preferences differ between the sexes. Three male and three female budgerigars were presented with four 8-s sound sequences in a preference apparatus: Simple (no pitch or temporal variation), Melody (pitch variation only), Rhythm (temporal variation only), and Complex (both pitch and temporal variation). Preference was quantified as the time spent in the area associated with each stimulus. No statistically significant differences among the four stimuli were detected within individuals. However, effect-size estimates indicated that two females spent more time with sequences containing rhythmic structure, whereas males showed no consistent preference related to either melodic or rhythmic components. Multidimensional scaling further suggested greater separation among stimulus conditions in females than in males. Consistent with this pattern, condition differentiation indices were higher in all three females than in all three males, although the sex difference was not statistically significant. These results suggest a possible sex-related difference in how budgerigars behaviorally weight temporal structure, with females showing greater differentiation among auditory sequence types under the present testing conditions.

## Introduction

Human auditory communication is organized along two major dimensions: a spectral dimension, including pitch and frequency structure, and a temporal dimension, including rhythm and interval structure. In spoken language, relatively slow changes in fundamental frequency contribute to prosody and intonation and are associated more strongly with right-hemisphere processing, whereas faster temporal patterns contribute to phonological grouping and syntactic processing and are associated more strongly with left-hemisphere mechanisms (Zatorre, Belin & Penhune et al., 2002; Poeppel, 2003; Oderbolz, Poeppel & Meyer, 2025). A broadly parallel distinction can be drawn in music, in which melody depends primarily on pitch contour, whereas rhythm and meter depend on temporal regularities. Although spectral and temporal information recruit partly distinct neural systems, they are integrated during language and music perception to support prediction, emotional expression, and structural processing (Peretz & Zatorre, 2005; Koelsch, 2011; Peretz, Vuvan, Lagrois & Armony, 2015).

This framework motivates comparative studies of auditory sequence perception in non-human animals. Examining how animals respond to melodic (pitch-based) and rhythmic (time-based) organization can help clarify the evolutionary origins of auditory processing abilities relevant to human language and music. Behavioral preference tests using sequences that isolate or combine melodic and rhythmic features provide one way to assess the relative weighting of these dimensions in auditory evaluation

Vocal-learning birds are particularly informative models because of their sophisticated auditory categorization abilities and several parallels with human speech acquisition (Doupe & Kuhl, 1999; Dooling & Okanoya, 1987). Among vocal learners, parrots such as the budgerigar (Melopsittacus undulatus) provide useful comparative models. Budgerigars show accurate pitch perception (Weisman et al., 2004), can detect complex harmonic changes (Lohr & Dooling, 1998), can discriminate human vowels (Dooling & Brown, 1990), and perform well in sequence-discrimination tasks, showing sensitivity to changes in the temporal ordering of elements within vocal sequences (Lawson et al., 2018; Fishbein et al., 2019). Vocal-auditory behavior in birds, however, often differs between the sexes. In many species, males sing more than females (Riebel, 2019), and sex differences in the rate and form of vocal learning have been reported even in species in which both sexes vocalize (Heil, Plummer, & Striedter, 2000).

In many songbirds, song learning is strongly male-biased, and females of some species show limited adult song production or learning. Budgerigars are notable because both males and females are vocal learners and share broadly similar vocal repertoires.

Female budgerigars show slower group-level call convergence, whereas males often exhibit rapid whole-call imitation (Heil & Striedter, 2000). Nevertheless, the capacity for adult vocal learning in both sexes makes this species useful for investigating sex-related differences in auditory evaluation without the pronounced production dimorphism found in many songbirds.

Previous studies have examined temporal and rhythmic processing in budgerigars using operant and motor tasks. Budgerigars can exhibit motor entrainment to regular auditory tempos, although with limited precision and after extensive training (Hasegawa et al., 2011), and their tapping behavior can be influenced by metronome distractors in a self-paced task (Seki & Tomyta, 2019). They are also sensitive to the temporal ordering of elements in species-natural (Tu & Dooling, 2012) and non-natural sequences (Spierings & ten Cate, 2016), as well as to acoustic cues associated with lexical stress (Hoeschele & Fitch, 2016). Importantly, sex differences in rhythmic preference have been reported: female budgerigars preferred stimuli with rhythmic structure, whereas males tended to prefer arrhythmic stimuli (Hoeschele & Bowling, 2016).

Budgerigars also show accurate pitch perception with low critical ratios across the frequency range relevant to their contact calls (Okanoya & Dooling, 1987; Dent, Dooling, & Pierce, 2000; Weisman et al., 2004), can detect complex harmonic changes (Lohr & Dooling, 1998), and can discriminate human vowels (Dooling & Brown, 1990), indicating high sensitivity to pitch variation. In contrast to the growing evidence for temporal and rhythmic sensitivity, however, the behavioral evaluation of melodic pitch patterns in budgerigars remains less well studied. Wagner et al. (2020) found no clear preference between two melodies in budgerigars; however, this experiment was designed to test sensitivity to consonance versus dissonance rather than to measure melodic preference itself, which may explain the absence of differential choice. Thus, it remains unclear whether budgerigars differentially weight temporal structure, pitch structure, or combinations of both when evaluating sound sequences. Given that both male and female zebra finches show strong sensitivity to pitch-based prosodic cues in human speech (Spierings & ten Cate, 2014), however, it is reasonable to infer that budgerigars—which possess even finer frequency discrimination—are likely to be sensitive to the presence or absence of melodic elements within acoustic sequences.

Natural budgerigar vocalizations include long, complex warble songs composed of many syllable types that vary in both temporal patterning and acoustic structure. Because these syllables differ in spectral properties, natural vocal sequences contain substantial pitch-based variation as well as time-based structure. Budgerigars therefore encounter both dimensions in their acoustic environment, making it plausible that both contribute to auditory sequence evaluation (Tu, Osmanski & Dooling, 2011).

The present study investigated behavioral responses to sound sequences that differed in melodic (pitch-based) and rhythmic (temporal) organization. I used a preference-based paradigm in which birds could move freely between areas associated with different auditory stimuli, and spontaneous preference was quantified as the time spent in each stimulus area. Preference measures do not directly establish perceptual discrimination, but they can provide indices of stimulus salience, behavioral weighting, and reinforcing value. Specifically, I asked whether (1) budgerigars show differential preferences for rhythmic or melodic structure and (2) these response patterns differ between males and females. By addressing these questions, the study aimed to clarify how a vocal-learning bird behaviorally weights spectral and temporal information in auditory sequences and to provide comparative data relevant to the processing of structured acoustic signals.

## Methods

### Subjects

Six adult budgerigars (*Melopsittacus undulatus*; three males and three females) obtained from a local breeder served as subjects. The birds were approximately one year old at the start of the experiment. Each bird was housed individually in a standard aviary cage (W 29 cm × D 37 cm × H 38 cm) under a 12:12 h light-dark cycle, with food and water available ad libitum. Before testing, the birds were acclimated to the experimental room for at least two weeks.

All procedures were approved by the Teikyo University Animal Care and Use Committee (permission no. 23-012) and were conducted in accordance with national guidelines for the care and use of laboratory animals.

### Apparatus

Behavioral preference was measured using a three-compartment apparatus with a stimulus area at each end. The two stimulus areas were constructed from cages of the same type as the birds’ home cages and were arranged at right angles to each other. A central neutral area, in which no auditory stimulus was presented, separated the two stimulus areas (Figure 1). This silent central area allowed time spent near an auditory stimulus to be distinguished from time spent elsewhere in the apparatus.

**Figure 1.**
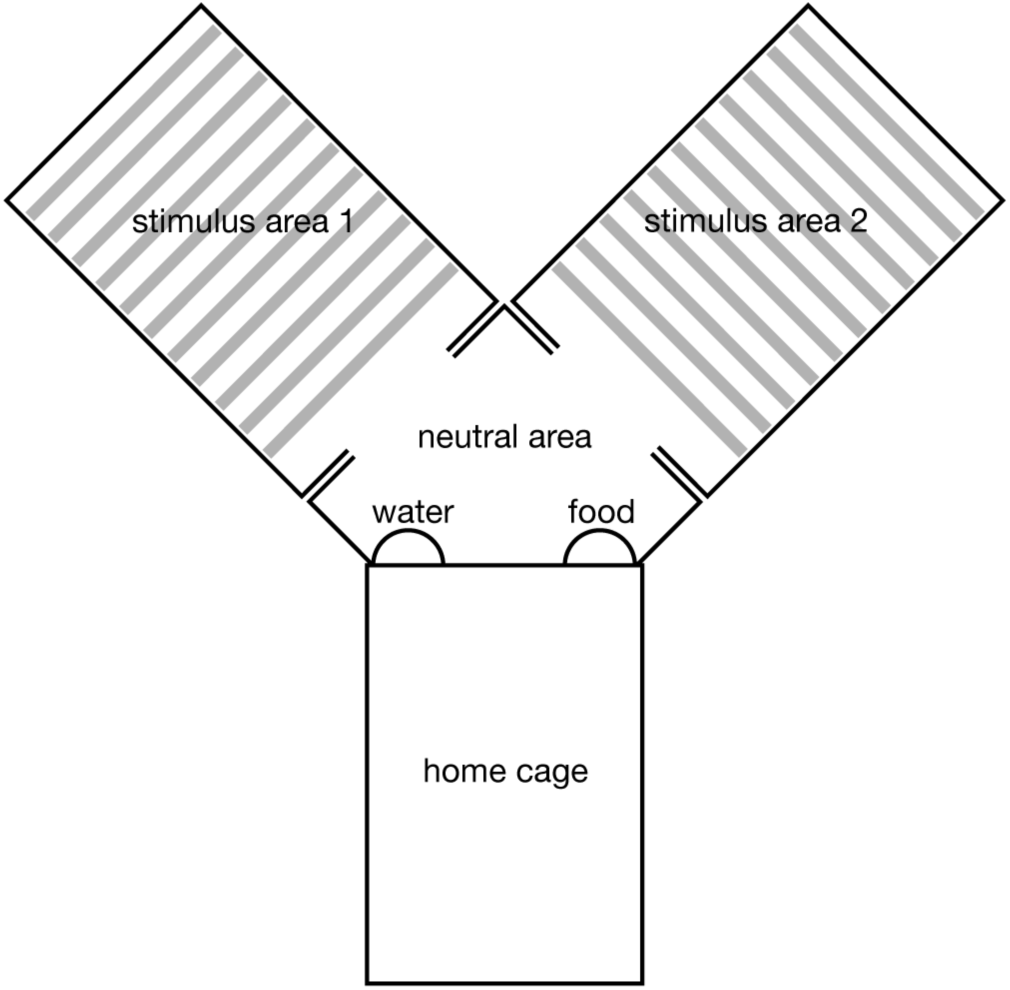
Schematic representation of the preference apparatus. The apparatus is shown from above. Two stimulus areas were arranged at right angles to each other and connected by a central neutral area. A loudspeaker was positioned beneath each stimulus area to present the auditory stimulus assigned to that side.

The floor of each stimulus area was equipped with perches spaced at 2-cm intervals. Food and water were available in the neutral area. During each session, birds were allowed to move freely throughout the apparatus. Bird position was recorded at 10 Hz using Arduino-connected weight sensors installed beneath the floor of each stimulus area; the sensors responded to loads exceeding approximately 5 g. When a bird entered and remained in either stimulus area, the stimulus assigned to that side was played through a loudspeaker (Bose Companion 2) concealed beneath the apparatus.

### Stimuli

The stimulus set was created by independently manipulating the presence or absence of rhythmic and melodic structure in the melody of Mary Had a Little Lamb. The original sequence contained notes ranging from C5 (523 Hz) to G5 (784 Hz), was presented at 120 beats per minute, and lasted 8 s.

The original sequence was modified to produce four stimuli: Simple, Melody, Rhythm, and Complex (Figure 2). The Simple sequence contained neither pitch variation nor the original temporal pattern and consisted of repeated eighth notes at 659 Hz (E5) at a constant tempo. The Melody sequence preserved the original pitch pattern but used only eighth notes at a constant tempo. The Rhythm sequence preserved the original temporal pattern but used a constant pitch of 659 Hz (E5). The Complex sequence preserved both the original melodic and rhythmic patterns. Each stimulus therefore consisted of an 8-s sound sequence.

**Figure 2.**
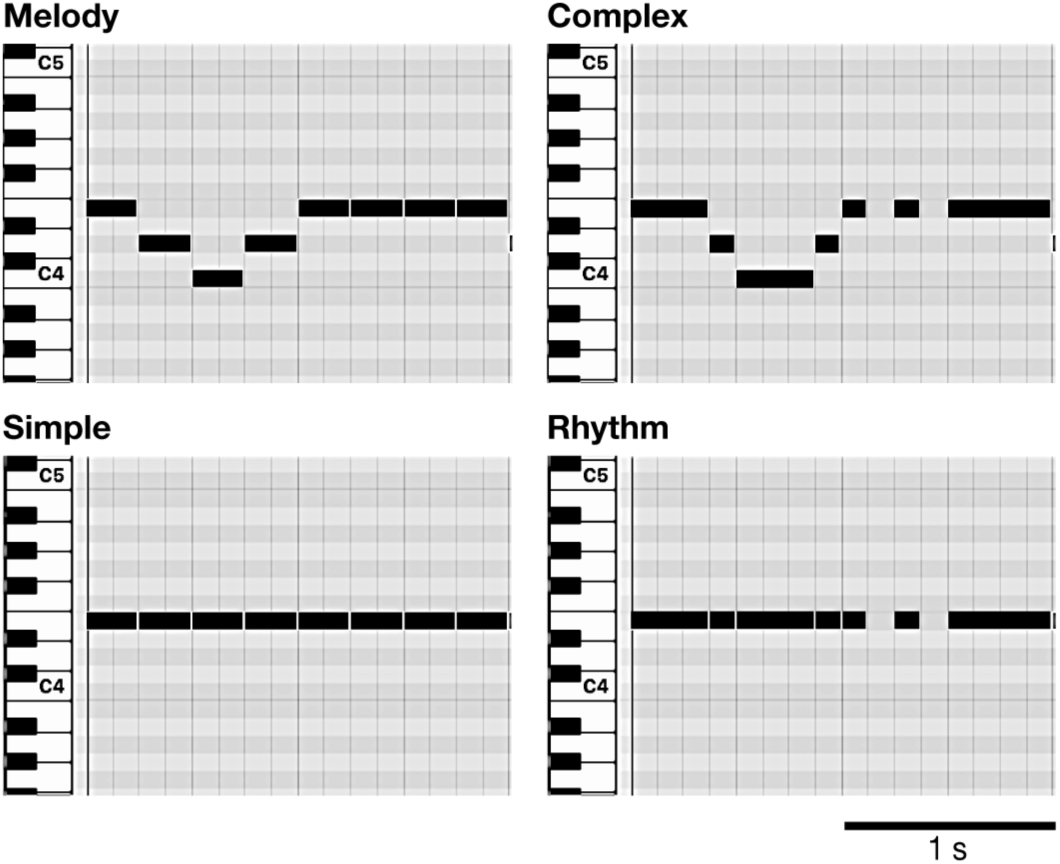
Piano-roll representations of the four auditory stimuli. The first 2 s of each 8-s stimulus are shown for the Simple, Melody, Rhythm, and Complex conditions. Horizontal position represents time, and vertical position represents pitch. The four stimuli differed in the presence or absence of melodic (pitch) and rhythmic (temporal) variation.

While a bird remained in a stimulus area, the stimulus assigned to that side was presented repeatedly, with a 2-s silent interval between repetitions. The playback level was approximately 70 dB(A), measured at the center of the stimulus area. Stimuli were generated using a whistle-like timbre (Korg Wavestation, JB-Whistle patch) and rendered in GarageBand (Apple Inc.).

### Procedure

Before each experimental session, the bird’s home cage was placed in front of the preference apparatus. The session began after the bird voluntarily entered the apparatus and the door of the home cage was closed.

Each session began with a 30-min habituation period during which no sounds were presented, followed by a 2-h stimulation phase. During the stimulation phase, entry into either stimulus area triggered playback of the stimulus assigned to that side, whereas the neutral area remained silent.

In each session, one of the six possible pairwise combinations of the four stimuli was selected. Each pair was tested twice with the stimulus-location assignment reversed, yielding 12 sessions per individual. Thus, each stimulus was presented in six sessions for each bird. The order of stimulus pairs was pseudorandomized across sessions within each individual.

Preference for each stimulus was quantified as the cumulative time spent in the corresponding stimulus area during the stimulation phase. To reduce habituation, each bird was tested once or twice per week.

Experiments were conducted in a vivarium to reduce stress and freezing associated with complete social isolation. The apparatus was surrounded by 50-mm-thick sound-absorbing panels (Tokyo Bouon, Japan), and visual contact with other birds was blocked by opaque plastic panels. Other cages in the room were also enclosed by sound-absorbing panels.

### Data Analysis

All statistical analyses were conducted using R (R Core Team, 2026) in RStudio (version 2026.05.1+225), with statistical significance defined as α = .05. Given the small sample size and the potential for non-normal distributions, nonparametric methods were used.

All four conditions were tested in each individual. However, because each session included only one pair drawn from the six possible pairwise combinations, responses to a given stimulus were obtained under different pairing contexts across sessions. The four conditions therefore could not be treated as fully matched observations within a single repeated-measures structure. Accordingly, nonparametric comparisons that did not assume paired observations were performed separately for each individual.

For each bird, a Kruskal-Wallis test was first used to assess differences in staying time among the four stimulus conditions. Dunn’s tests were then used for post hoc pairwise comparisons, with Holm correction for multiple comparisons. Cliff’s δ was calculated as a nonparametric measure of effect size. Analyses were conducted separately for each bird because multiple sessions were obtained from the same individual and pooling sessions across birds would have introduced pseudoreplication.

To examine the contributions of melodic and rhythmic components more directly, data from relevant conditions were pooled. Conditions containing melodic structure (Melody+: Melody and Complex) were compared with conditions lacking melodic structure (Melody−: Simple and Rhythm). Similarly, conditions containing rhythmic structure (Rhythm+: Rhythm and Complex) were compared with conditions lacking rhythmic structure (Rhythm−: Simple and Melody). These pooled contrasts were analyzed within each individual using two-sided Mann-Whitney U tests, with Cliff’s δ used to estimate effect size.

In addition to the within-individual preference analyses, the degree of behavioral differentiation among stimulus conditions was compared across birds.

Multidimensional scaling (MDS) was used to visualize the relative similarity of responses to the four stimulus conditions. Staying-time data were first z-standardized within each individual to reduce the influence of differences in overall activity level. Pairwise distances between staying-time distributions were then calculated using the Wasserstein distance, which captures differences between distributions rather than differences in central tendency alone.

For each individual, the resulting distance matrix was projected into a two-dimensional MDS space. Conditions located closer together were interpreted as producing more similar distributions of staying time, whereas greater separation indicated greater behavioral differentiation between conditions.

To quantify overall differentiation among the four stimulus conditions, a condition differentiation index was calculated for each individual as the mean of the six pairwise Wasserstein distances. Larger values indicate greater differentiation among stimulus-related staying-time distributions, whereas smaller values indicate more similar responses across conditions.

## Results

### Behavioral responses across stimulus conditions

Staying-time distributions for the four stimulus conditions are shown for each bird in Figure 3. The degree of overlap among conditions varied across individuals: some birds showed apparent differences in median staying time, whereas others showed substantial overlap among distributions.

**Figure 3.**
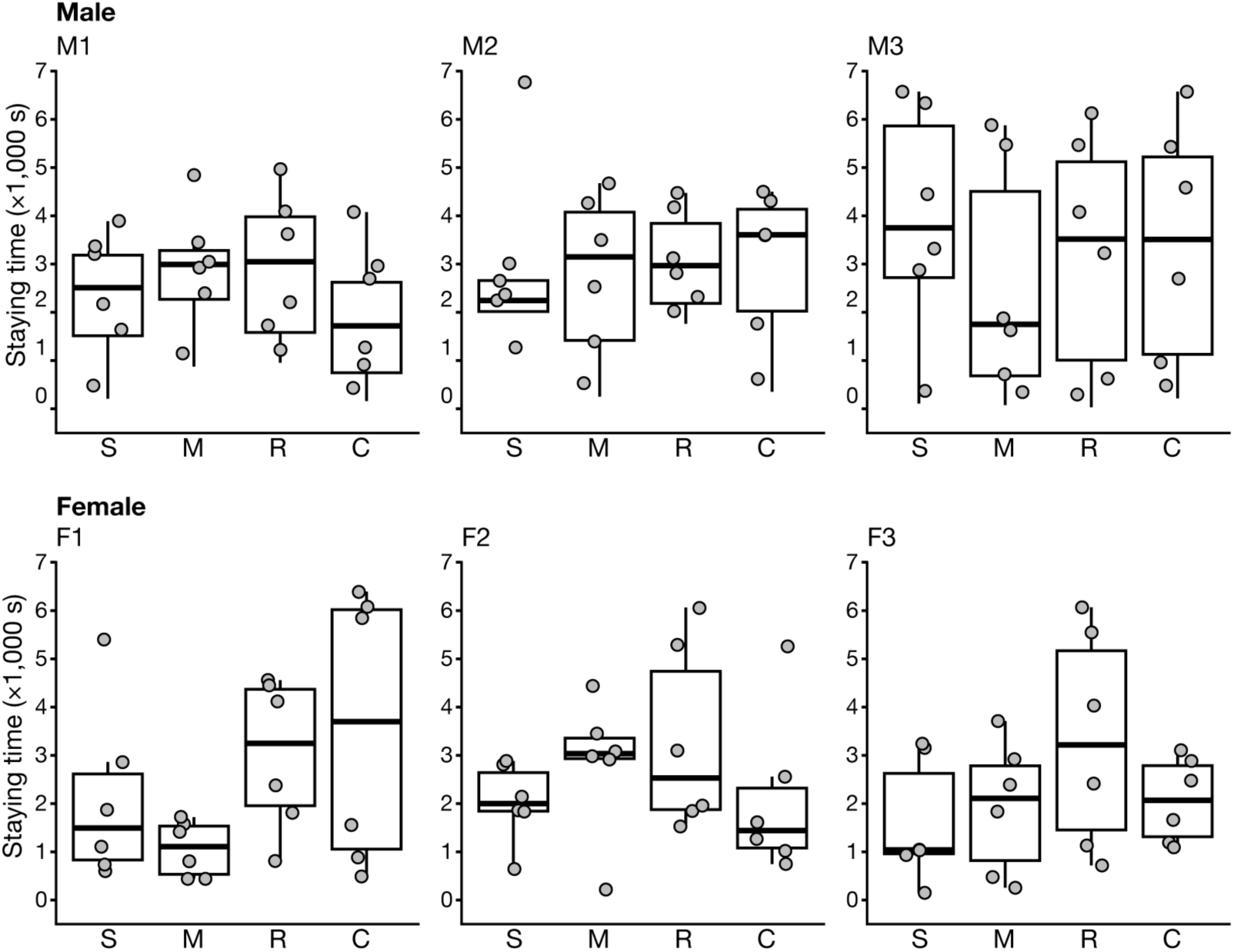
Staying time for the four auditory stimulus conditions in each budgerigar. Distributions of cumulative staying time are shown separately for the Simple (S), Melody (M), Rhythm **(R),** and Complex (C) conditions for each individual. Boxplots show the median and interquartile range, and individual observations are shown as jittered points. Upper and lower panels represent male and female individuals, respectively.

Kruskal-Wallis tests conducted separately for each individual detected no statistically significant effect of stimulus condition. Subsequent Dunn’s pairwise comparisons with Holm correction likewise yielded no statistically significant differences. Nevertheless, several pairwise contrasts showed medium-to-large Cliff’s δ values, indicating potentially meaningful differences in response magnitude that warrant cautious consideration.

For F1, staying time was longer for the Rhythm than for the Simple sequence (δ = 0.333). The Melody sequence was associated with shorter staying time than the Simple (δ = 0.389), Rhythm (δ = 0.833), and Complex (δ = 0.500) sequences. For F2, staying time was longer for the Melody sequence than for the Simple (δ = 0.792), Rhythm (δ = 0.792), and Complex (δ = 0.333) sequences; the Rhythm sequence also elicited longer staying time than the Complex sequence (δ = 0.556). For F3, staying time was longer for the Rhythm sequence than for the Simple (δ = 0.720), Melody (δ = 0.611), and Complex (δ = 0.444) sequences. For M1, the Complex sequence was associated with shorter staying time than the Simple (δ = 0.333), Melody (δ = 0.333), and Rhythm (δ = 0.444) sequences. M2 and M3 showed little differentiation among the four conditions, including in the effect-size estimates.

To assess the contributions of melodic and rhythmic structure more directly, Melody+ versus Melody− and Rhythm+ versus Rhythm− conditions were compared within each individual (Figure 4). A statistically significant difference was detected only for the Rhythm+ versus Rhythm− contrast in F1 (U = 37, p = 0.046). Effect-size estimates nevertheless suggested longer staying times in Rhythm+ than in Rhythm− conditions for F1 (δ = 0.486) and F3 (δ = 0.333). For these birds, the corresponding Melody+ versus Melody− effects were smaller (F1, δ = 0.208; F3, δ = 0.056). F2 showed some differentiation among the four individual stimulus conditions, but little evidence that this pattern was attributable to the melody or rhythm dimension alone (Rhythm+ vs. Rhythm−, δ = 0.056; Melody+ vs. Melody−, δ = 0.042). None of the three males showed a consistent preference associated with either melodic or rhythmic structure in these pooled contrasts.

**Figure 4.**
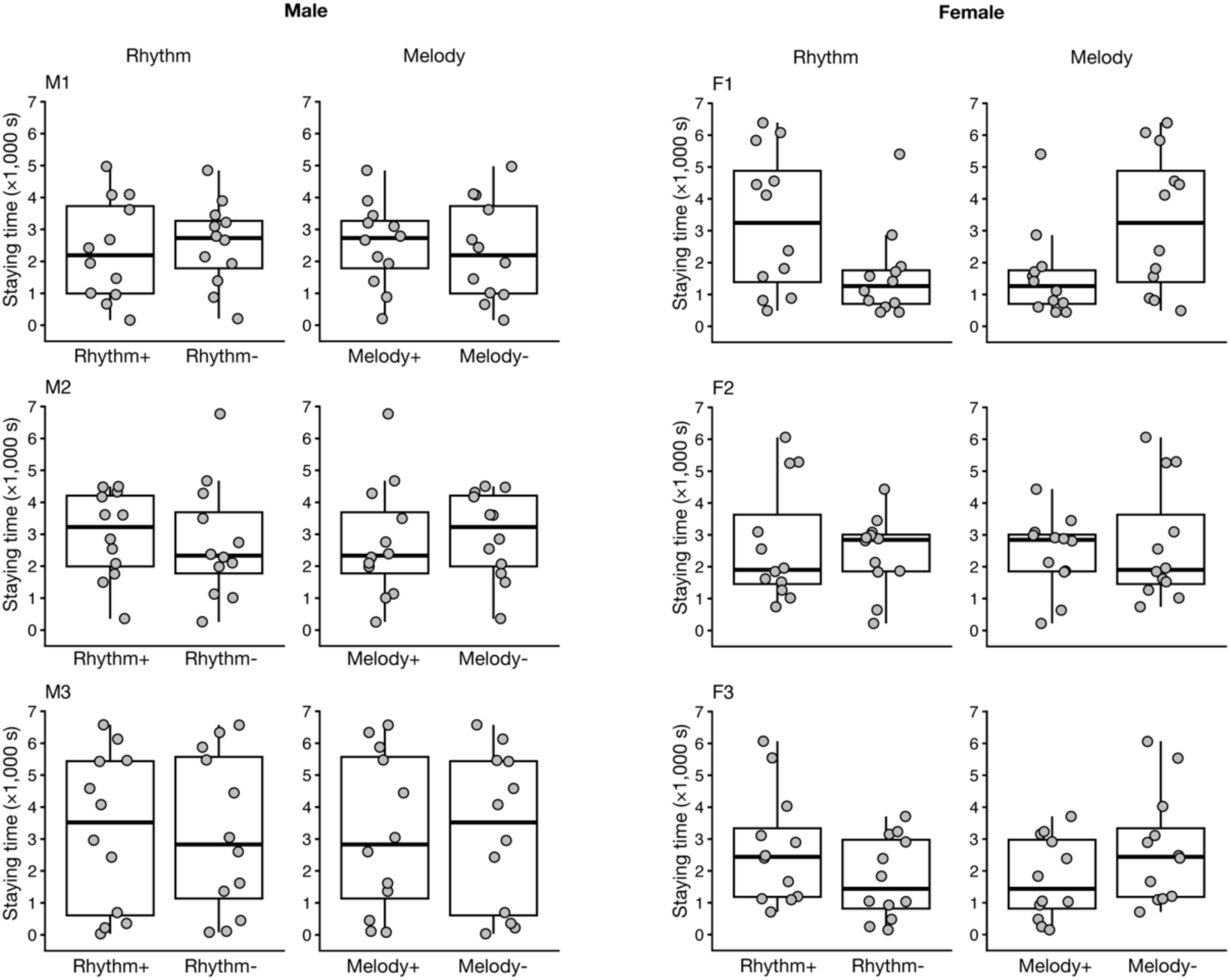
Effects of melodic and rhythmic structure on staying time in each budgerigar. Staying times were pooled according to the presence or absence of rhythmic structure (Rhythm+ vs. Rhythm-} and melodic structure (Melody+ vs. Melody-) and compared separately within each individual. Boxplots show the median and interquartile range, and individual observations are shown as jittered points. The left and right halves represent male and female individuals, respectively.

MDS was used to compare the relative differentiation of responses among the four stimulus conditions across birds (Figure 5). Visual inspection suggested greater separation among conditions in females, whereas the conditions tended to cluster more closely in males. To summarize this descriptive pattern, overall differentiation was quantified using the condition differentiation index.

**Figure 5.**
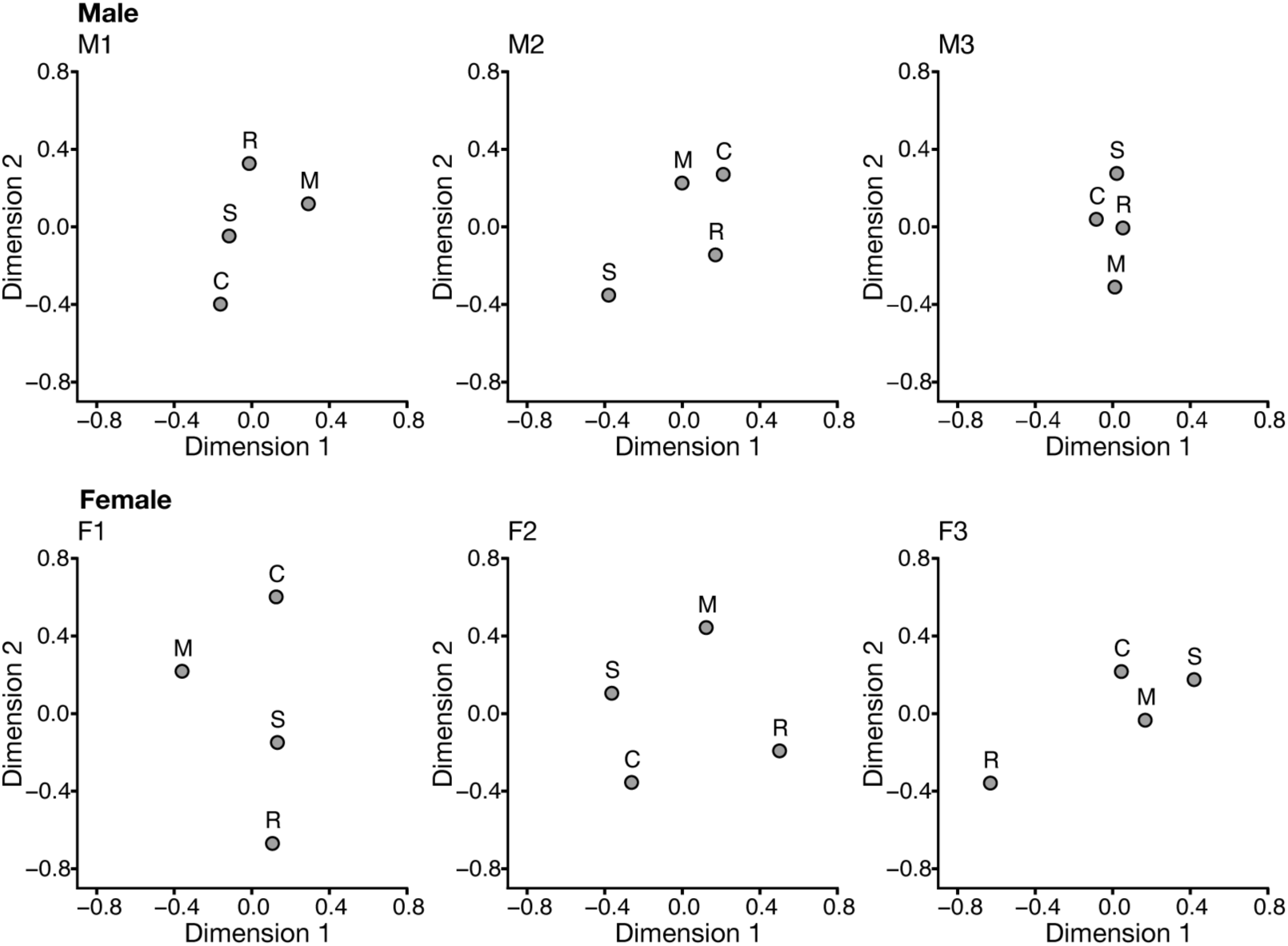
Differentiation of behavioral responses among the four auditory stimulus conditions. Multidimensional scaling (MOS) representations were derived from pairwise Wasserstein distances between z-standardized staying-time distributions for the Simple (S), Melody (M), Rhythm (R), and Complex (C) conditions in each individual. Greater separation between conditions indicates greater differentiation in behavioral responses. Upper and lower panels represent male and female individuals, respectively.

Condition differentiation indices were higher in each female than in any male (F1, F2, and F3: 0.796, 0.740, and 0.659, respectively; M1, M2, and M3: 0.499, 0.533, and 0.352, respectively). A Mann-Whitney U test did not detect a statistically significant sex difference (U = 9, p = 0.10). However, the estimated effect size was large (Cliff’s δ = 1.0), reflecting complete separation of the observed index values between the three females and three males in this sample.

## Discussion

The present study examined how budgerigars behaviorally evaluate auditory sequences that differ in melodic (spectral) and rhythmic (temporal) structure, and whether these response patterns differ between the sexes. Most inferential tests yielded no statistically significant differences. Nevertheless, effect-size estimates and individual-level analyses revealed several recurring patterns that may be informative for understanding auditory sequence evaluation in this species.

Overall, the results do not support a single dominant acoustic cue that uniformly determined preference across birds. Instead, there was substantial inter-**individual** variation in the relative weighting of melodic and rhythmic structure. Two females (F1 and F3) showed effect-size patterns favoring rhythm-containing conditions, whereas F2 showed condition-specific differences that were not readily attributable to melody or rhythm alone. These **individual** differences suggest that the behavioral value of acoustic structure may vary considerably among budgerigars. Such individual variation in the degree of abstraction derived from auditory stimuli has also been reported previously (ten Cate et al., 2016). In a rhythm discrimination study, all three female budgerigars successfully learned to distinguish the regular from the irregular rhythm during training; however, they differed markedly in their ability to generalize this discrimination to tempo-altered test stimuli, with only one individual showing sensitivity to global regularity.

A previous study reported a sex difference in rhythmic preference in budgerigars, with females preferring **rhythmic** stimuli and males preferring **arrhythmic** stimuli (Hoeschele & Bowling, 2016). In the present study, two of the three females showed a tendency to spend more time with stimuli containing rhythmic structure, whereas none of the males showed a comparable pattern. The present experiment examined responses to the presence or absence of rhythmic and melodic elements within acoustic sequences rather than to the regularity or irregularity of these elements. Therefore, male budgerigars might have shown differential preferences if the regularity of rhythmic structure had been manipulated. Although the present sample is too small to establish a general sex difference, the convergence between these findings raises the possibility that regular temporal structure may have particular behavioral salience for at least some female budgerigars.

Notably, preferences associated with either melodic or rhythmic structure did not necessarily translate into the strongest preference for the Complex sequence, which combined both features. Thus, within the present preference paradigm, the effects of melodic and rhythmic structure were not simply additive. This pattern is compatible with the possibility that spectral and temporal information can contribute partly independently to behavioral evaluation, although the present data do not establish independent perceptual processing mechanisms. This finding differs from accounts of human auditory cognition in which prosodic and syntactic information can interact during hierarchical processing (Cutler et al., 1997), but direct cross-species comparisons should be made cautiously because the present stimuli and behavioral measures differ substantially from those used in human language studies.

The sex-related pattern in overall condition differentiation also warrants consideration. Although the between-sex comparison was not statistically significant, all three females had higher condition differentiation indices than all three males, yielding a large effect-size estimate. The MDS representations were similarly more dispersed in females than in males. These observations suggest greater behavioral differentiation among stimulus types in the females tested here. One possible interpretation is that females evaluate acoustic variation more selectively, perhaps because discrimination among vocal features can have ecological relevance in social or mate-choice contexts (Riebel, 2019). This interpretation, however, remains tentative given the small number of birds.

In contrast, males showed more clustered MDS representations and relatively uniform preference responses across stimulus types. This pattern is unlikely, by itself, to indicate an inability to discriminate the stimuli, given the well-documented auditory sensitivity of budgerigars. One possible explanation is that the present stimuli were encountered in a passive setting without an explicit social context. Vocal production learning in budgerigars is strongly influenced by social interaction, and auditory exposure alone can be less effective than exposure accompanied by direct interaction with conspecifics (Farabaugh et al., 1994; Osmanski et al., 2021). Accordingly, the weak differentiation observed in males may reflect similar motivational or social values assigned to all four stimuli under the present conditions rather than poor perceptual discrimination.

Sex-specific responses to socially meaningful vocal signals have also been reported in other parrots. Male and female orange-fronted conures respond differently to convergent and divergent contact-call interactions, indicating that the behavioral significance of acoustic variation can depend on the sex of the receiver (Balsby & Scarl, 2008). In light of these findings, the present results raise the testable possibility that female budgerigars may show relatively differentiated responses to acoustic structure even when explicit social cues are absent, whereas males’ preference responses may depend more strongly on the social context in which a signal is presented. Direct manipulation of social context will be required to evaluate this possibility.

Several limitations related to sample size should be considered when interpreting these results. First, the study included only three birds of each sex, which limits statistical power for between-sex comparisons and the extent to which the observed sex-related patterns can be generalized to the population level. The medium-to-large effect sizes observed in several comparisons should therefore be regarded as estimates of potentially meaningful patterns rather than as evidence of established population-level effects. Replication with a larger number of individuals will be necessary to determine the reliability and generality of these patterns.

A related limitation concerns the number of repeated preference measurements obtained within each individual. Because repeated exposure to the stimuli might have reduced overall activity through habituation, the number of measurements for each stimulus pair was intentionally limited to no more than four. This restriction may have reduced statistical power for detecting within-individual differences in preference. Increasing the number of measurements per stimulus pair, while controlling for possible habituation, might have provided more stable estimates of individual preference and allowed firmer conclusions regarding differences among stimulus conditions.

Second, preference-based paradigms assess behavioral valuation rather than perceptual discrimination directly. A reliable preference indicates that stimuli differ in their behavioral consequences for the subject, but the absence of a preference does not imply that the stimuli are perceptually indistinguishable. Perceptually distinct stimuli may elicit similar staying times if they have comparable motivational value. The present results should therefore be interpreted primarily as differences in behavioral evaluation or attentional weighting of auditory patterns rather than as definitive differences in perceptual resolution.

Future studies combining preference tests with operant discrimination and generalization tasks should help distinguish behavioral valuation from perceptual sensitivity. In particular, systematic manipulation of spectral and temporal cues during generalization tests could provide a more direct assessment of how each dimension contributes to sequence discrimination. Experiments that vary the social context of stimulus presentation would also test whether the sex-related patterns observed here depend on the social relevance of acoustic signals. Finally, examining neural correlates of these behaviors may help determine whether sex-related variation reflects differences in auditory processing, attention, or motivational systems.

In conclusion, the present preference-based study provides preliminary evidence that melodic and rhythmic structure may be weighted differently across individual budgerigars and suggests a possible sex-related pattern in auditory sequence evaluation. Two of the three females showed response patterns favoring rhythm-containing stimuli, and all females showed greater overall differentiation among stimulus conditions than did the males, although most inferential tests, including the between-sex comparison, were not statistically significant. The relatively uniform responses of males may reflect the limited motivational or social relevance of passively presented stimuli rather than an inability to discriminate among them. Taken together, these findings identify rhythmic structure and social context as promising factors for further investigation and support the value of combining preference and discrimination measures with socially contextualized testing paradigms to clarify how male and female budgerigars evaluate complex auditory sequences.

## Supporting information

Summary of Data

R script for across-stimulus analysis

R script for subjective distance across-stimuli

## Acknowledgement

This work was supported by JSPS KAKENHI Grant Number JP21K12606 to MW.

## References

Balsby, T.J., Scarl, J.C., (2008). Sex-specific responses to vocal convergence and divergence of contact calls in orange-fronted conures (*Aratinga canicularis*). Proc. Biol. Sci. 275 (1647), 2147–2154. doi: 10.1098/rspb.2008.0517.

Cutler, A., Dahan, D., van Donselaar, W. (1997). Prosody in the comprehension of spoken language: A literature review. Language and Speech, 40 (2), 141–201. 10.1177/002383099704000203.

Dent, M.L., Dooling, R.J., Pierce, A.S. (2000) Frequency discrimination in budgerigars (Melopsittacus undulatus): effects of tone duration and tonal context. J. Acoust. Soc. Am., 107 (5 Pt 1), 2657–2664. doi: 10.1121/1.428651.

Dooling, R.J., Brown, S.D. (1990). Speech perception by budgerigars (*Melopsittacus undulatus*): Spoken vowels. Percept. Psychophys., 47 (6), 568– 574. 10.3758/BF03203109.

Doupe, A.J., Kuhl, P.K., (1992) Birdsong and human speech: common themes and mechanisms. Annu. Rev. Neurosci., 22, 567–631. doi: 10.1146/annurev.neuro.22.1.567.

Farabaugh, S.M., Linzenbold, A., Dooling, R.J., (1994) Vocal plasticity in budgerigars (*Melopsittacus undulatus*): evidence for social factors in the learning of contact calls. J. Comp. Psychol., 108 (1), 81–92. doi: 10.1037/0735-7036.108.1.81.

Fishbein, A.R., Idsardi, W.J., Ball, G.F., Dooling, R.J., (2019) Sound sequences in birdsong: how much do birds really care? Philos. Trans. R. Soc. Lond. B Biol. Sci., 375 (1789), 20190044. doi: 10.1098/rstb.2019.0044.

Hasegawa, A., Okanoya, K., Hasegawa, T., Seki, Y., (2011) Rhythmic synchronization tapping to an audio-visual metronome in budgerigars. Sci. Rep., 1, 120. doi: 10.1038/srep00120.

Hile, A.G., Plummer, T.K., Striedter, G.F., (2000) Male vocal imitation produces call convergence during pair bonding in budgerigars, *Melopsittacus undulatus*. Anim. Behav., 59 (6), 1209–1218. doi: 10.1006/anbe.1999.1438.

Hile, A.G., Striedter, G.F., (2000) Call Convergence within Groups of Female Budgerigars (*Melopsittacus undulatus*). Ethology, 106, 1105–1114. 10.1046/j.1439-0310.2000.00637.x.

Hoeschele, M., Bowling, D.L., (2016) Sex Differences in Rhythmic Preferences in the Budgerigar (*Melopsittacus undulatus*): A Comparative Study with Humans. Front. Psychol., 7, 1543. doi: 10.3389/fpsyg.2016.01543.

Hoeschele, M., Fitch, W.T., (2016) Phonological perception by birds: budgerigars can perceive lexical stress. Anim. Cogn., 19 (3), 643–654. doi: 10.1007/s10071-016-0968-3.

Koelsch, S., (2011) Toward a Neural Basis of Music Perception – A Review and Updated Model. Front. Psychology, 2, 110. doi: 10.3389/fpsyg.2011.00110.

Lawson, S.L., Fishbein, A.R., Prior, N.H., Ball, G.F., Dooling, R.J., (2018) Relative salience of syllable structure and syllable order in zebra finch song. Anim. Cogn. 21 (4), 467–480. doi: 10.1007/s10071-018-1182-2.

Lohr, B., Dooling, R.J., (1998) Detection of changes in timbre and harmonicity in complex sounds by zebra finches (*Taeniopygia guttata*) and budgerigars (*Melopsittacus undulatus*). J. Comp. Psychol., 112 (1), 36–47. doi: 10.1037/0735-7036.112.1.36.

Oderbolz, C., Poeppel, D., Meyer, M., (2025). Asymmetric Sampling in Time: Evidence and perspectives. Neurosci. Biobehav. Rev., 171, 1–17. 10.1016/j.neubiorev.2025.106082.

Okanoya, K., Dooling, R.J., (1987). Hearing in passerine and psittacine birds: A comparative study of absolute and masked auditory thresholds. J. Comp. Psychol., 101 (1), 7–15. 10.1037/0735-7036.101.1.7.

Osmanski, M.S., Seki, Y., Dooling, R.J., (2021) Constraints on vocal production learning in budgerigars (*Melopsittacus undulates*). Learn. Behav., 49 (1), 150–158. doi: 10.3758/s13420-021-00465-6.

Peretz, I., Vuvan, D., Lagrois, M-É., Armony, J.L., (2015) Neural overlap in processing music and speech. Philos. Trans. R. Soc. Lond. B Biol. Sci., 370, 20140090. 10.1098/rstb.2014.0090.

Peretz, I., Zatorre, R.J., (2005) Brain organization for music processing. Annu. Rev. Psychol., 56, 89–114. doi: 10.1146/annurev.psych.56.091103.070225.

Poeppel, D., (2003) The analysis of speech in different temporal integration windows: cerebral lateralization as ‘asymmetric sampling in time’, Speech Commun., 41 (1), 245–255. 10.1016/S0167-6393(02)00107-3.

Riebel, K., Odom, K.J., Langmore, N.E., Hall, M.L., (2019) New insights from female bird song: towards an integrated approach to studying male and female communication roles. Biol. Lett., 15, 20190059. 10.1098/rsbl.2019.0059.

Seki, Y., Tomyta, K., (2019). Effects of metronomic sounds on a self-paced tapping task in budgerigars and humans. Curr. Zool., 65, 121–128. doi: 10.1093/cz/zoy075.

Spierings, M.J., ten Cate, C., (2014). Zebra finches are sensitive to prosodic features of human speech. Proc. Biol. Sci., 281, 20140480. doi: 10.1098/rspb.2014.0480.

Spierings, M.J., Ten Cate, C., (2016) Budgerigars and zebra finches differ in how they generalize in an artificial grammar learning experiment. Proc. Natl. Acad. Sci. U. S. A. 113 (27), E3977–84. doi: 10.1073/pnas.1600483113.

ten Cate, C., Spierings, M., Hubert, J., Honing, H., (2016) Can Birds Perceive Rhythmic Patterns? A Review and Experiments on a Songbird and a Parrot Species. Front. Psychol., 7, 730. doi: 10.3389/fpsyg.2016.00730.

Tu, H.W., Dooling, R.J., (2012) Perception of warble song in budgerigars (*Melopsittacus undulatus*): evidence for special processing. Anim. Cogn., 15 (6), 1151–1159. doi: 10.1007/s10071-012-0539-1.

Tu, H.W., Osmanski, M.S., Dooling, R.J., (2011) Learned vocalizations in budgerigars (*Melopsittacus undulatus*): the relationship between contact calls and warble song. J. Acoust. Soc. Am. 129 (4), 2289–97. doi: 10.1121/1.3557035.

Wagner, B., Bowling, D.L., Hoeschele, M., (2020) Is consonance attractive to budgerigars? No evidence from a place preference study. Anim. Cogn., 23 (5), 973–987. doi: 10.1007/s10071-020-01404-0.

Weisman, R.G., Njegovan, M.G., Williams, M.T., Cohen, J.S., Sturdy, C.B., (2004) A behavior analysis of absolute pitch: sex, experience, and species. Behav. Process., 66, 289–307. doi: 10.1016/j.beproc.2004.03.010.

Zatorre, R.J., Belin, P., Penhune, V.B., (2002) Structure and function of auditory cortex: music and speech. Trends Cogn. Sci., 6 (1), 37–46. doi: 10.1016/s1364-6613(00)01816-7.

