## Supplementary material for "Exploring rhythmic and melodic preferences in budgerigars: Individual and possible sex-related variation": R script for across-stimulus analysis

#========================================================
# R script for nonparametric statistical analyses
#========================================================
#
# This script performs the following analyses:
#
# 1. Kruskal-Wallis test comparing staying times among
#    the four stimulus conditions.
#
# 2. Calculation of Cliff's delta for all six pairwise
#    comparisons among the four stimulus conditions.
#
# 3. Mann-Whitney U tests and Cliff's delta comparing:
#    (a) rhythm-present vs. rhythm-absent conditions, and
#    (b) melody-present vs. melody-absent conditions.
#
# Raw staying-time data were used for all analyses.
#
# Stimulus conditions:
# C1: melody absent, rhythm absent
# C2: melody present, rhythm absent
# C3: melody absent, rhythm present
# C4: melody present, rhythm present

#========================================================
# 1. Enter staying-time data
#========================================================

# Six observations were obtained for each stimulus condition.
# F1 data is shown below as a sample dataset.

C1_NoM_NoR <- c(
  18752, 53972, 11106, 28616, 6044, 7375
)

C2_YesM_NoR <- c(
  15723, 4441, 14132, 8024, 17193, 4442
)

C3_NoM_YesR <- c(
  8134, 45575, 23738, 18123, 44534, 41193
)

C4_YesM_YesR <- c(
  60798, 63897, 4904, 8867, 58385, 15558
)

#========================================================
# 2. Kruskal-Wallis test across the four stimulus conditions
#========================================================

# The Kruskal-Wallis test was used to examine whether
# staying times differed among the four stimulus conditions
# within the individual.

staying_time <- c(
  C1_NoM_NoR,
  C2_YesM_NoR,
  C3_NoM_YesR,
  C4_YesM_YesR
)

condition <- factor(
  rep(
    c(
      "C1_NoM_NoR",
      "C2_YesM_NoR",
      "C3_NoM_YesR",
      "C4_YesM_YesR"
    ),
    each = 6
  )
)

data_kw <- data.frame(
  staying_time = staying_time,
  condition = condition
)

kw_result <- kruskal.test(
  staying_time ~ condition,
  data = data_kw
)

# Display the Kruskal-Wallis test result.
kw_result

#========================================================
# 3. Define a function for Cliff's delta
#========================================================

# Cliff's delta was calculated as:
# delta = [number of (x > y) pairs - number of (x < y) pairs]
#         / total number of x-y pairs
#
# Delta ranges from -1 to +1.
#
# A positive value indicates that observations in the first
# condition tend to be larger than those in the second condition.
# A negative value indicates that observations in the first
# condition tend to be smaller than those in the second condition.

cliffs_delta <- function(x, y) {
  
  comparisons <- outer(
    x,
    y,
    FUN = "-"
  )
  
  delta <- (
    sum(comparisons > 0) -
      sum(comparisons < 0)
  ) / length(comparisons)
  
  return(delta)
}

#========================================================
# 4. Cliff's delta for all pairwise comparisons
#========================================================

# Cliff's delta was calculated for all six possible pairwise
# comparisons among the four stimulus conditions:
#
# C1 vs. C2
# C1 vs. C3
# C1 vs. C4
# C2 vs. C3
# C2 vs. C4
# C3 vs. C4

delta_C1_C2 <- cliffs_delta(
  C1_NoM_NoR,
  C2_YesM_NoR
)

delta_C1_C3 <- cliffs_delta(
  C1_NoM_NoR,
  C3_NoM_YesR
)

delta_C1_C4 <- cliffs_delta(
  C1_NoM_NoR,
  C4_YesM_YesR
)

delta_C2_C3 <- cliffs_delta(
  C2_YesM_NoR,
  C3_NoM_YesR
)

delta_C2_C4 <- cliffs_delta(
  C2_YesM_NoR,
  C4_YesM_YesR
)

delta_C3_C4 <- cliffs_delta(
  C3_NoM_YesR,
  C4_YesM_YesR
)

#========================================================
# 5. Classify the magnitude of Cliff's delta
#========================================================

# Effect-size magnitude was classified using the following
# commonly used thresholds:
#
# |delta| < 0.147          : negligible
# 0.147 <= |delta| < 0.330 : small
# 0.330 <= |delta| < 0.474 : medium
# |delta| >= 0.474         : large
#
# These categories describe effect-size magnitude only and
# should not be interpreted as significance tests.

classify_delta <- function(delta) {
  
  abs_delta <- abs(delta)
  
  if (abs_delta < 0.147) {
    return("negligible")
    
  } else if (abs_delta < 0.330) {
    return("small")
    
  } else if (abs_delta < 0.474) {
    return("medium")
    
  } else {
    return("large")
  }
}

# Combine all pairwise Cliff's delta values into a table.

cliff_pairwise_results <- data.frame(
  
  Comparison = c(
    "C1 vs C2",
    "C1 vs C3",
    "C1 vs C4",
    "C2 vs C3",
    "C2 vs C4",
    "C3 vs C4"
  ),
  
  Cliffs_delta = c(
    delta_C1_C2,
    delta_C1_C3,
    delta_C1_C4,
    delta_C2_C3,
    delta_C2_C4,
    delta_C3_C4
  )
)

cliff_pairwise_results$Magnitude <- sapply(
  cliff_pairwise_results$Cliffs_delta,
  classify_delta
)

# Display all pairwise effect sizes.
cliff_pairwise_results

#========================================================
# 6. Comparison of rhythm-present and rhythm-absent conditions
#========================================================

# Rhythm-present conditions:
# C3 (melody absent, rhythm present)
# C4 (melody present, rhythm present)
#
# Rhythm-absent conditions:
# C1 (melody absent, rhythm absent)
# C2 (melody present, rhythm absent)
#
# The six observations from each relevant stimulus condition
# were pooled, resulting in 12 observations for each level
# of the rhythm factor.

Rhythm_present <- c(
  C3_NoM_YesR,
  C4_YesM_YesR
)

Rhythm_absent <- c(
  C1_NoM_NoR,
  C2_YesM_NoR
)

# Perform a two-sided Mann-Whitney U test.
#
# R's wilcox.test() returns the Wilcoxon rank-sum statistic W.
# The corresponding Mann-Whitney U statistic is calculated below.

rhythm_test <- wilcox.test(
  Rhythm_present,
  Rhythm_absent,
  alternative = "two.sided",
  exact = FALSE
)

n1_rhythm <- length(Rhythm_present)

rhythm_U <- unname(rhythm_test$statistic) -
  n1_rhythm * (n1_rhythm + 1) / 2


# Calculate Cliff's delta.
#
# A positive delta indicates that staying times tend to be
# longer in the rhythm-present conditions.
# A negative delta indicates that staying times tend to be
# longer in the rhythm-absent conditions.

delta_rhythm <- cliffs_delta(
  Rhythm_present,
  Rhythm_absent
)

magnitude_rhythm <- classify_delta(delta_rhythm)

#========================================================
# 7. Comparison of melody-present and melody-absent conditions
#========================================================

# Melody-present conditions:
# C2 (melody present, rhythm absent)
# C4 (melody present, rhythm present)
#
# Melody-absent conditions:
# C1 (melody absent, rhythm absent)
# C3 (melody absent, rhythm present)
#
# The six observations from each relevant stimulus condition
# were pooled, resulting in 12 observations for each level
# of the melody factor.

Melody_present <- c(
  C2_YesM_NoR,
  C4_YesM_YesR
)

Melody_absent <- c(
  C1_NoM_NoR,
  C3_NoM_YesR
)

# Perform a two-sided Mann-Whitney U test.

melody_test <- wilcox.test(
  Melody_present,
  Melody_absent,
  alternative = "two.sided",
  exact = FALSE
)

n1_melody <- length(Melody_present)

melody_U <- unname(melody_test$statistic) -
  n1_melody * (n1_melody + 1) / 2


# Calculate Cliff's delta.
#
# A positive delta indicates that staying times tend to be
# longer in the melody-present conditions.
# A negative delta indicates that staying times tend to be
# longer in the melody-absent conditions.

delta_melody <- cliffs_delta(
  Melody_present,
  Melody_absent
)

magnitude_melody <- classify_delta(delta_melody)


#========================================================
# 8. Summary table for rhythm and melody comparisons
#========================================================

# The results of the collapsed rhythm and melody comparisons
# are summarized below.
#
# Positive Cliff's delta values indicate longer staying times
# in the feature-present condition (Rhythm+ or Melody+).

collapsed_results <- data.frame(
  
  Comparison = c(
    "Rhythm present vs Rhythm absent",
    "Melody present vs Melody absent"
  ),
  
  U = c(
    rhythm_U,
    melody_U
  ),
  
  p_value = c(
    rhythm_test$p.value,
    melody_test$p.value
  ),
  
  Cliffs_delta = c(
    delta_rhythm,
    delta_melody
  ),
  
  Magnitude = c(
    magnitude_rhythm,
    magnitude_melody
  )
)

# Display the summary table.
collapsed_results# R script for nonparametric statistical analyses
#========================================================
#
# This script performs the following analyses:
#
# 1. Kruskal-Wallis test comparing staying times among
#    the four stimulus conditions.
#
# 2. Calculation of Cliff's delta for all six pairwise
#    comparisons among the four stimulus conditions.
#
# 3. Mann-Whitney U tests and Cliff's delta comparing:
#    (a) rhythm-present vs. rhythm-absent conditions, and
#    (b) melody-present vs. melody-absent conditions.
#
# Raw staying-time data were used for all analyses.
#
# Stimulus conditions:
# C1: melody absent, rhythm absent
# C2: melody present, rhythm absent
# C3: melody absent, rhythm present
# C4: melody present, rhythm present

#========================================================
# 1. Enter staying-time data
#========================================================

# Six observations were obtained for each stimulus condition.
# F1 data is shown below as a sample dataset.

C1_NoM_NoR <- c(
  18752, 53972, 11106, 28616, 6044, 7375
)

C2_YesM_NoR <- c(
  15723, 4441, 14132, 8024, 17193, 4442
)

C3_NoM_YesR <- c(
  8134, 45575, 23738, 18123, 44534, 41193
)

C4_YesM_YesR <- c(
  60798, 63897, 4904, 8867, 58385, 15558
)

#========================================================
# 2. Kruskal-Wallis test across the four stimulus conditions
#========================================================

# The Kruskal-Wallis test was used to examine whether
# staying times differed among the four stimulus conditions
# within the individual.

staying_time <- c(
  C1_NoM_NoR,
  C2_YesM_NoR,
  C3_NoM_YesR,
  C4_YesM_YesR
)

condition <- factor(
  rep(
    c(
      "C1_NoM_NoR",
      "C2_YesM_NoR",
      "C3_NoM_YesR",
      "C4_YesM_YesR"
    ),
    each = 6
  )
)

data_kw <- data.frame(
  staying_time = staying_time,
  condition = condition
)

kw_result <- kruskal.test(
  staying_time ~ condition,
  data = data_kw
)

# Display the Kruskal-Wallis test result.
kw_result

#========================================================
# 3. Define a function for Cliff's delta
#========================================================

# Cliff's delta was calculated as:
#
# delta = [number of (x > y) pairs - number of (x < y) pairs]
#         / total number of x-y pairs
#
# Delta ranges from -1 to +1.
#
# A positive value indicates that observations in the first
# condition tend to be larger than those in the second condition.
# A negative value indicates that observations in the first
# condition tend to be smaller than those in the second condition.

cliffs_delta <- function(x, y) {
  
  comparisons <- outer(
    x,
    y,
    FUN = "-"
  )
  
  delta <- (
    sum(comparisons > 0) -
      sum(comparisons < 0)
  ) / length(comparisons)
  
  return(delta)
}

#========================================================
# 4. Cliff's delta for all pairwise comparisons
#========================================================

# Cliff's delta was calculated for all six possible pairwise
# comparisons among the four stimulus conditions:
#
# C1 vs. C2
# C1 vs. C3
# C1 vs. C4
# C2 vs. C3
# C2 vs. C4
# C3 vs. C4

delta_C1_C2 <- cliffs_delta(
  C1_NoM_NoR,
  C2_YesM_NoR
)

delta_C1_C3 <- cliffs_delta(
  C1_NoM_NoR,
  C3_NoM_YesR
)

delta_C1_C4 <- cliffs_delta(
  C1_NoM_NoR,
  C4_YesM_YesR
)

delta_C2_C3 <- cliffs_delta(
  C2_YesM_NoR,
  C3_NoM_YesR
)

delta_C2_C4 <- cliffs_delta(
  C2_YesM_NoR,
  C4_YesM_YesR
)

delta_C3_C4 <- cliffs_delta(
  C3_NoM_YesR,
  C4_YesM_YesR
)

#========================================================
# 5. Classify the magnitude of Cliff's delta
#========================================================

# Effect-size magnitude was classified using the following
# commonly used thresholds:
#
# |delta| < 0.147          : negligible
# 0.147 <= |delta| < 0.330 : small
# 0.330 <= |delta| < 0.474 : medium
# |delta| >= 0.474         : large
#
# These categories describe effect-size magnitude only and
# should not be interpreted as significance tests.

classify_delta <- function(delta) {
  
  abs_delta <- abs(delta)
  
  if (abs_delta < 0.147) {
    return("negligible")
    
  } else if (abs_delta < 0.330) {
    return("small")
    
  } else if (abs_delta < 0.474) {
    return("medium")
    
  } else {
    return("large")
  }
}

# Combine all pairwise Cliff's delta values into a table.

cliff_pairwise_results <- data.frame(
  
  Comparison = c(
    "C1 vs C2",
    "C1 vs C3",
    "C1 vs C4",
    "C2 vs C3",
    "C2 vs C4",
    "C3 vs C4"
  ),
  
  Cliffs_delta = c(
    delta_C1_C2,
    delta_C1_C3,
    delta_C1_C4,
    delta_C2_C3,
    delta_C2_C4,
    delta_C3_C4
  )
)

cliff_pairwise_results$Magnitude <- sapply(
  cliff_pairwise_results$Cliffs_delta,
  classify_delta
)

# Display all pairwise effect sizes.
cliff_pairwise_results

#========================================================
# 6. Comparison of rhythm-present and rhythm-absent conditions
#========================================================

# Rhythm-present conditions:
# C3 (melody absent, rhythm present)
# C4 (melody present, rhythm present)
#
# Rhythm-absent conditions:
# C1 (melody absent, rhythm absent)
# C2 (melody present, rhythm absent)
#
# The six observations from each relevant stimulus condition
# were pooled, resulting in 12 observations for each level
# of the rhythm factor.

Rhythm_present <- c(
  C3_NoM_YesR,
  C4_YesM_YesR
)

Rhythm_absent <- c(
  C1_NoM_NoR,
  C2_YesM_NoR
)

# Perform a two-sided Mann-Whitney U test.
#
# R's wilcox.test() returns the Wilcoxon rank-sum statistic W.
# The corresponding Mann-Whitney U statistic is calculated below.

rhythm_test <- wilcox.test(
  Rhythm_present,
  Rhythm_absent,
  alternative = "two.sided",
  exact = FALSE
)

n1_rhythm <- length(Rhythm_present)

rhythm_U <- unname(rhythm_test$statistic) -
  n1_rhythm * (n1_rhythm + 1) / 2

# Calculate Cliff's delta.
#
# A positive delta indicates that staying times tend to be
# longer in the rhythm-present conditions.
# A negative delta indicates that staying times tend to be
# longer in the rhythm-absent conditions.

delta_rhythm <- cliffs_delta(
  Rhythm_present,
  Rhythm_absent
)

magnitude_rhythm <- classify_delta(delta_rhythm)

#========================================================
# 7. Comparison of melody-present and melody-absent conditions
#========================================================

# Melody-present conditions:
# C2 (melody present, rhythm absent)
# C4 (melody present, rhythm present)
#
# Melody-absent conditions:
# C1 (melody absent, rhythm absent)
# C3 (melody absent, rhythm present)
#
# The six observations from each relevant stimulus condition
# were pooled, resulting in 12 observations for each level
# of the melody factor.

Melody_present <- c(
  C2_YesM_NoR,
  C4_YesM_YesR
)

Melody_absent <- c(
  C1_NoM_NoR,
  C3_NoM_YesR
)

# Perform a two-sided Mann-Whitney U test.

melody_test <- wilcox.test(
  Melody_present,
  Melody_absent,
  alternative = "two.sided",
  exact = FALSE
)

n1_melody <- length(Melody_present)

melody_U <- unname(melody_test$statistic) -
  n1_melody * (n1_melody + 1) / 2

# Calculate Cliff's delta.
#
# A positive delta indicates that staying times tend to be
# longer in the melody-present conditions.
# A negative delta indicates that staying times tend to be
# longer in the melody-absent conditions.

delta_melody <- cliffs_delta(
  Melody_present,
  Melody_absent
)

magnitude_melody <- classify_delta(delta_melody)

#========================================================
# 8. Summary table for rhythm and melody comparisons
#========================================================

# The results of the collapsed rhythm and melody comparisons
# are summarized below.
#
# Positive Cliff's delta values indicate longer staying times
# in the feature-present condition (Rhythm+ or Melody+).

collapsed_results <- data.frame(
  
  Comparison = c(
    "Rhythm present vs Rhythm absent",
    "Melody present vs Melody absent"
  ),
  
  U = c(
    rhythm_U,
    melody_U
  ),
  
  p_value = c(
    rhythm_test$p.value,
    melody_test$p.value
  ),
  
  Cliffs_delta = c(
    delta_rhythm,
    delta_melody
  ),
  
  Magnitude = c(
    magnitude_rhythm,
    magnitude_melody
  )
)

# Display the summary table.
collapsed_results
