## Supplementary material for "Exploring rhythmic and melodic preferences in budgerigars: Individual and possible sex-related variation": R script for subjective distance across-stimuli

# ======================================================
# R script for MDS analysis and Condition Differentiation Index
# ======================================================

# This script performs the following analyses:
# 1. Within-individual z-standardization of staying-time data
# 2. Calculation of pairwise Wasserstein distances
# 3. Calculation of the Condition Differentiation Index (CDI)
# 4. Multidimensional scaling (MDS) based on Wasserstein distances
#
# C1: melody absent, rhythm absent
# C2: melody present, rhythm absent
# C3: melody absent, rhythm present
# C4: melody present, rhythm present

# ======================================================
# 1. Enter staying-time data
# ======================================================

# Six observations were obtained for each stimulus condition.
# F1 data is shown below as a sample dataset.

C1_NoM_NoR <- c(
  18752, 53972, 11106, 28616, 6044, 7375
)
C2_YesM_NoR <- c(
  15723, 4441, 14132, 8024, 17193, 4442
)
C3_NoM_YesR <- c(
  8134, 45575, 23738, 18123, 44534, 41193
)
C4_YesM_YesR <- c(
  60798, 63897, 4904, 8867, 58385, 15558
)

# ======================================================
# 2. Combine the four stimulus conditions into a list
# ======================================================

data_raw <- list(
  C1_NoM_NoR   = C1_NoM_NoR,
  C2_YesM_NoR  = C2_YesM_NoR,
  C3_NoM_YesR  = C3_NoM_YesR,
  C4_YesM_YesR = C4_YesM_YesR
)

# ======================================================
# 3. Within-individual z-standardization
# ======================================================

# All 24 staying-time observations (4 conditions x 6 observations)
# were pooled within the individual.
# The mean and standard deviation were calculated from these
# 24 observations, and each observation was converted to a z score.
#
# This standardization was used to reduce the influence of
# individual differences in the overall scale of staying time
# and to facilitate comparison of relative preference patterns
# across individuals.

all_values <- unlist(data_raw)
mean_all <- mean(all_values)
sd_all   <- sd(all_values)

# Convert raw staying times to z scores.
z_all <- (all_values - mean_all) / sd_all

# Split the standardized values back into the four conditions.
data_z <- split(
  z_all,
  rep(names(data_raw), each = 6)
)

# Display the standardized data.
data_z

# ======================================================
# 4. Define a function for the Wasserstein distance
# ======================================================

# Because all four conditions contain the same number of observations
# (n = 6), the one-dimensional Wasserstein distance can be calculated
# as the mean absolute difference between the corresponding ordered
# observations of the two distributions.

wasserstein_distance <- function(x, y) {
  mean(abs(sort(x) - sort(y)))
}

# ======================================================
# 5. Calculate pairwise Wasserstein distances
# ======================================================

# A 4 x 4 dissimilarity matrix was constructed from the pairwise
# Wasserstein distances among the four stimulus conditions.

condition_names <- names(data_z)
distance_matrix <- matrix(
  0,
  nrow = 4,
  ncol = 4,
  dimnames = list(condition_names, condition_names)
)

for (i in 1:4) {
  
  for (j in 1:4) {
    
    distance_matrix[i, j] <-
      wasserstein_distance(
        data_z[[i]],
        data_z[[j]]
      )
  }
}

# Display the Wasserstein distance matrix.
round(distance_matrix, 3)

# ======================================================
# 6. Calculate the Condition Differentiation Index (CDI)
# ======================================================

# With four stimulus conditions, there are six unique pairwise
# Wasserstein distances.
#
# The CDI was defined as the mean of these six distances.
# A larger CDI indicates greater behavioral differentiation
# among the four stimulus conditions.

distance_object <- as.dist(distance_matrix)
differentiation_index <- mean(distance_object)

# Display the CDI.
differentiation_index

# ======================================================
# 7. Perform multidimensional scaling (MDS)
# ======================================================

# Classical multidimensional scaling was applied to the
# Wasserstein distance matrix to represent the relative
# relationships among the four stimulus conditions
# in a two-dimensional space.
#
# The orientation and sign of the MDS dimensions are arbitrary.
# Therefore, interpretation should be based on the relative
# distances among stimulus conditions rather than on the
# direction or sign of the MDS axes.

mds_result <- cmdscale(
  distance_object,
  k = 2,
  eig = TRUE
)

# Extract the two-dimensional MDS coordinates.
mds_coordinates <- as.data.frame(mds_result$points)
colnames(mds_coordinates) <- c("MDS1", "MDS2")
mds_coordinates$Condition <- rownames(mds_coordinates)

# Display the MDS coordinates.
mds_coordinates

# ======================================================
# 8. Plot the MDS configuration
# ======================================================

# Conditions located closer together represent more similar
# staying-time distributions, whereas conditions located farther
# apart represent greater behavioral differentiation.

plot(
  mds_coordinates$MDS1,
  mds_coordinates$MDS2,
  type = "n",
  xlab = "MDS Dimension 1",
  ylab = "MDS Dimension 2",
  main = "MDS of stimulus conditions"
)

points(
  mds_coordinates$MDS1,
  mds_coordinates$MDS2,
  pch = 21,
  bg = "gray",
  cex = 1.5
)

text(
  mds_coordinates$MDS1,
  mds_coordinates$MDS2,
  labels = mds_coordinates$Condition,
  pos = 3,
  cex = 0.8
)

# ======================================================
# 9. Export the MDS coordinates
# ======================================================

write.csv(
  mds_coordinates,
  "MDS_coordinates.csv",
  row.names = FALSE
)

# ======================================================
# 10. Export the Wasserstein distance matrix
# ======================================================

write.csv(
  distance_matrix,
  "Wasserstein_distance_matrix.csv"
)
